# The aging whole blood transcriptome reveals a potential role of FASLG in COVID-19

**DOI:** 10.1101/2020.12.04.412494

**Authors:** Luiz Gustavo de Almeida Chuffa, Jeferson dos Santos Souza, Mariana Costa de Mello, Mario de Oliveira Neto, Robson Francisco Carvalho

## Abstract

The risk for severe illness from COVID-19 increases with age as older patients are at the highest risk. Although it is still unclear whether the virus is blood-transmitted, the viral RNA is detected in serum. Identifying how Severe Acute Respiratory Syndrome Coronavirus 2 (SARS-CoV-2) interacts with specific blood components during aging is expected to guide proper therapies. Considering that all human coronavirus require host cellular molecules to promote infection, we investigated the aging whole blood transcriptome from the Genotype-Tissue Expression (GTEx) database to explore differentially expressed genes (DEGs) translated into proteins potentially interacting with viral proteins. From a total of 22 DEGs in aged blood, five genes (*FASLG, CTSW, CTSE, VCAM1*, and *BAG3*) changed expression during aging. These age-related genes are involved in immune response, inflammation, cell component and cell adhesion, and platelet activation/aggregation. Both males and females older than 50 overexpress *FASLG* compared with younger adults (20-30 years old), possibly inducing a hyper-inflammatory cascade that activates specific immune cells. Furthermore, the expression of cathepsins (*CTSW* and *CTSE*) and the anti-apoptotic co-chaperone molecule *BAG3* was significantly increased throughout aging in both gender. By exploring publicly available Single-Cell RNA-Sequencing (scRNA-Seq) data on peripheral blood of SARS-CoV-2-infected patients, we found *FASLG* and *CTSW* expressed mainly in natural killer (NK) cells and CD8+ (cytotoxic) T lymphocytes whereas *BAG3* was expressed in CD4+ T cells, naive T cells, and CD14+ monocytes. The increased expression of *FASLG* in blood during aging may explain why older patients are more prone to severe acute viral infection complications. These results indicate *FASLG* as a prognostic candidate and potential therapeutic target for more aggressive clinical manifestation of COVID-19.

## Introduction

Coronavirus disease 2019 (COVID-19) is a pandemic infection caused by severe acute respiratory syndrome coronavirus 2 (SARS-CoV-2) [1]. This virus is associated with a broad spectrum of respiratory disturbances, varying from upper airway symptoms to aggressive pneumonia [2]. In lung parenchyma, it produces alveolar edema, fibrin deposition, and hemorrhage [2]. Notably, vascular changes are one of the distinctive features of COVID-19. Many patients have demonstrated clinical signs of thrombotic microangiopathy [3], with intravascular coagulation and thrombosis associated with multisystem organ failure [4].

Although SARS-CoV-2 affects the lungs primordially, there is convincing evidence that it alters the coagulation processes in severe cases [5]. The formation of blood clots disrupts circulation due to thrombosis, pulmonary embolism, and heart attacks [3, 4]. The association of these changes in coagulation and protein aggregates with alterations in inflammatory parameters results in an increase of COVID-19 mortality at alarming proportions. The micro-thrombotic environment arises from hyperactivation of the coagulation cascade associated with hyper-intense inflammation and immune activities [6]. Of note, the clotting process is complex and likely orchestrated by the massive release of pro-inflammatory mediators, cytokines, and tumor necrosis factor (TNF), mainly released from monocytes and endothelial cells [6]. The fact that the mortality rate of patients aged 60 years and over is higher than those under 60 years is indisputable [7]. Most of these critical cases are associated with the “cytokine storm,” as these exacerbated immune reactions may lead to an early death of elderly people regardless of comorbidities related to more severe cases [8].

Recent findings are correlating the blood subtype with viral susceptibility and infection [9, 10]. However, the individual molecular machinery of blood cells may be detrimental for the grade and type of response, such as exacerbated or reduced inflammation. Considering that aging is one of the most significant risk factors for severe cases of COVID-19, it is essential to determine blood host genetic variation throughout the aging process. The evolutionary conservation between the 2019 novel SARS-CoV-2 and SARS-CoV [11] allows the possibility to understand similarities and differences between these coronaviruses into public databases. Using computational predictions of SARS-CoV–human protein-protein interactions (PPIs), we can identify possible mechanisms behind the viral infection and identify potential drug targets [12, 13].

Considering that older individuals, with or without comorbidities, are more prone to develop more severe cases of COVID-19, including those related to blood perturbations, we investigated the whole blood transcriptome data during aging using the Genotype-Tissue Expression (GTEx) database [14, 15]. This strategy provided significant insights into age-associated target genes and how they can predict SARS-CoV-2-interactions in aged blood components.

## Materials and Methods

### Whole blood transcriptome during aging

We used whole blood RNA-Seq data from 670 males and females available at the GTEx portal (release V8) (https://www.gtexportal.org/) [16]. The BioJupies platform (https://amp.pharm.mssm.edu/biojupies/) [17] was used to find the differentially expressed genes (DEGs) in whole blood samples over aging (20-79 years-old). The samples were selected and matched per age range: 30-39, 40-49, 50-59; 60-69 and 70-79. Then, age ranges were individually compared with young adults (aged 20-29). Genes with Log2 fold-change ≥ |1| ≤ |1| and false discovery rate (FDR) < 0.05 were considered as DEGs (Supplementary Tables 1-5). The DEGs were used to identify protein-protein interaction networks and perform gene ontology enrichment analyses (Supplementary Figure 1).

### Virus-host protein-protein interactions (PPIs) overlapping DEGs in whole blood with SARS-CoV-related perturbations

We first selected SARS-CoV-2 cell entry mediators using the human proteins available in the COVID-19 Cell Atlas (https://www.covid19cellatlas.org/). In addition to this screening, we compared upregulated and downregulated DEGs in the whole blood during aging with the corresponding proteins that interact with HCoVs (Supplementary Table 9) by using publicly available databases. To uncover HCoVs-human PPIs, we used data from the Pathogen–Host Interactome Prediction using Structure Similarity (P-HIPSTer, http://phipster.org/) database. P-HIPSTer is a broad protein catalog of the virus-human interactions upon structural information with an experimental validation rate of approximately 76% [18]. Our research has used P-HIPSTer to identify proteins potentially interacting with SARS-CoV-2 in other tissues [19, 20].

The relative expression of *FASLG, CTSW, CTSE, VCAM1, and BAG3* (TMM normalized; V8 cohort) was also performed independently in male and female samples using One-way ANOVA followed by Tukey’s test. The results were analyzed with GraphPad Prism v. 6.00 for Windows (GraphPad Software, La Jolla, California, USA). Significant differences were set at P < 0.05.

### Tissue-specific interaction networks of potential SARS-CoV-host blood interactome

The HumanBase tool (https://hb.flatironinstitute.org) [21] was used to provide whole blood-specific networks based on upregulated genes during aging that code for proteins potentially interacting with SARS-CoV according to P-HIPSTer. We considered five functional modules: co-expression, interaction, transcription factor binding, gene set enrichment analysis perturbations and microRNA targets as active interaction sources, applying a minimum interaction confidence of 0.12. The maximum number of genes was set as 15.

### Gene Ontology Enrichment analysis of differentially expressed genes during aging

We performed the Kyoto Encyclopedia of Genes and Genomes (KEGG) pathway enrichment analysis and Gene Ontology enrichment analysis (Biological Processes) to identify the functions of DEGs by using the EnrichR database (http://amp.pharm.mssm.edu/Enrichr/) [22]. Top enriched terms were generated according to the lowest p-value < 0.05 (Fisher’s exact test). The molecular function and protein class related to the blood components during aging were performed in the PANTHER classification system v. 11.0 (http://www.pantherdb.org/) [23]. We used the UniProtKB database (http://www.uniprot.org/) to obtain functional information of the identified proteins.

### Protein-protein interaction (PPI) networks based on blood gene expression profile during aging

The genes that appeared overexpressed in aged blood samples were analyzed by STRING online tool (https://string-db.org/) [24]. The metasearch STRING database (Search Tool for Retrieval of Interacting Genes, v. 10.5) was used for mapping PPI enrichment. We considered the following settings: text mining, databases, experiments, and co-occurrence as sources of interaction. The minimum interaction score was 0.900 (highest confidence); in the networks, the disconnected nodes were hidden to show reliable interactions exclusively. The PPI enrichment P-value indicates the statistical significance registered by STRING. (access in October 2020).

### Single-cell transcriptomic analysis of human peripheral blood cells

We investigated the expression of selected genes (*FASLG, CTSW, CTSE, VCAM* and *BAG3*) in distinct blood cell populations based on previously published human single-cell RNA-seq data [25]. This single-cell dataset is available at the COVID-19 Cell Atlas (https://www.covid19cellatlas.org/) and was explored using the cellxgene interactive viewer (https://cellxgene.cziscience.com). The dataset includes peripheral blood mononuclear cells (PBMCs) populations from patients hospitalized with COVID-19 (n=7); patients with acute respiratory distress syndrome (n = 4), and healthy controls displaying no diasease (n = 6).

### Structural analysis of the Fas and FasL

The genes Fas (genbank code: XP_006717882.1) and FasL (genbank code: NP_000630.1) were obtained on NCBI GenBank, and protein structure blast of these sequences were performed using NCBI blastp suite [26]. Next, PDB structures and FASTA sequences were downloaded on protein data Bank [27], and the sequences obtained from GenBank and PDB were aligned on LALIGN [28]. Finally, sequence molecular mass calculation was evaluated using ProtParam expasy [29].

### Data representation and analysis

The comparison of blood candidate genes was based on Venn diagrams using the Venny2.0 tool (https://bioinfogp.cnb.csic.es/tools/venny/index.html) [30]. Heatmaps and Scatter plots for clustering analyses were performed using the web tool Morpheus (https://software.broadinstitute.org/morpheus) [31]. Metascape was used to provide GO terms enrichment [32] obtained from aged blood genes that potentially interact with SARS-CoV-2.

## Results

### The number of differentially expressed genes and the complexity of associated functions increase in the whole blood during aging

The number of DEGs varied significantly throughout aging (Log2 FC ≥ |1| ≤ |1| and FDR < 0.05; Tables S1-S5). We detected an increased number of DEGs in subjects with 50-59 years old (62 DEGs; 53 up- and 9 downregulated, respectively), 60-69 (251 DEGs; 212 up- and 39 downregulated, respectively), being the older individuals of 70-79 with the highest number (913 DEGs; 498 up- and 415 downregulated, respectively). These DEGs were predicted into PPI networks associated with immune response, cytokine and receptor activity, defense response, response to a stimulus, and signaling receptor binding (Figure S1 A-C). Forty-one upregulated genes shared the transcriptional profile during aging and are associated with cytokine-cytokine receptor interaction, lymphocyte and natural killer chemotaxis, and eosinophil migration (FDR < 0.05, combined score > 86.06; Figure S1 D-E and Tables S6-S7). The most prominent molecular functions were binding, regulation, and catalytic activity, while protein classes included those related to immunity, transcriptional regulator, signaling molecule, and protein modifying enzyme (Figure S1 F). Seven down-regulated genes shared the transcriptional profile during aging and mostly associated with apolipoprotein receptor binding, regulation of low-density lipoprotein, protein autoprocessing, negative regulation of receptor binding, and receptor-mediated endocytosis (FDR < 0.05, combined score > 75.90; Figure S1 G-H and Tables S6-S8). The catalytic activity, molecular regulator, transducer, and transcription activities were the most common molecular functions, while protein classes included transcriptional regulator, protein binding activity, protein modifying enzyme, and carrier protein (Figure S1 I). Upregulated and downregulated profiles of DEGs are depicted in the volcano plots (Figure S1 J).

### Virus-host PPI interactions reveal increased expression of potential targets in whole blood during aging

Next, we used the list of DEGs during aging to identify transcripts translated into proteins that potentially interact with SARS-CoV proteins based on P-HIPSTer database. This analysis identified 22 genes with an age-dependent expression profile (Table S9; Figure 1 A). Younger individuals (20-29, 30-39, and 40-49 years old) showed similarities and were distinctly clustered than older individuals (50-59, 60-69, and 70-79 years old). By comparing clusters two by two, the age range of 20-29 years old showed a unique mean expression pattern compared to other aging groups (Figure S2). Among DEGs translated into predicted proteins interacting with SARS-CoV, most of them appeared overexpressed in the age range of 50-79 years old (Figure 1 B and C); while *FASLG* appeared overexpressed in individuals aged 50-59, 60-69, and 70-79, the *CTSW, CTSE, VCAM1 and BAG3* targets were commonly overexpressed in the two older groups of individuals (60-69 and 70-79 years-old). The exclusive DEGs in individuals of 60-79 years old were *HBQ1, HSPA5, EPB41L3, SRC, PDCD1, CD7, CD8A, IGSF9, EPB41L4A, FRMPD3, and SCN2B*. After investigating tissue-specific functional modules using these DEGs, we observed a consistent interaction of whole blood proteins as being mostly associated with natural killer cells (Figure 1 D).

**Figure 1.**
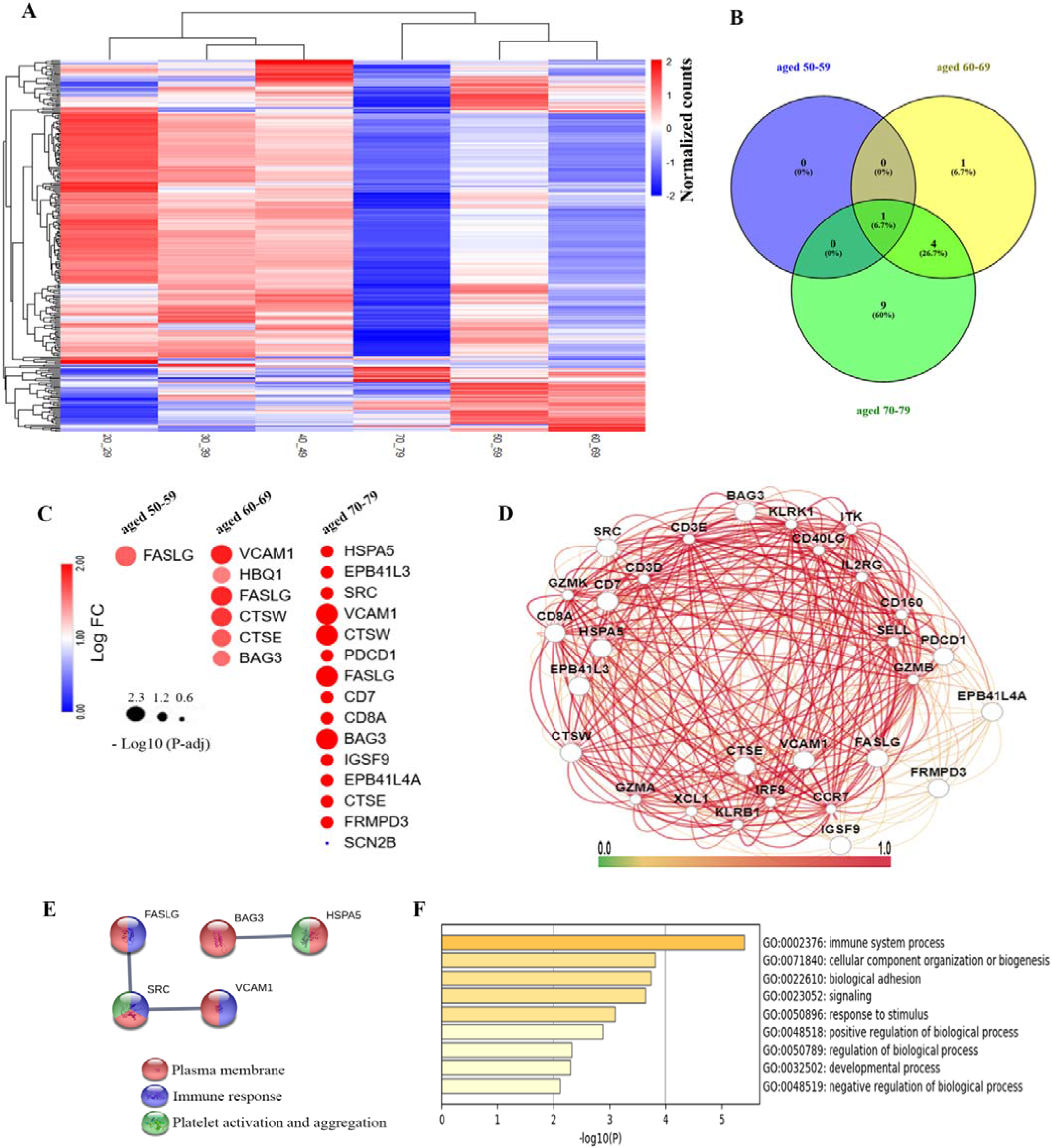
Identification of molecular targets that potentially interact with SARS-CoV-2 and its functions associated with whole blood during aging. A) Heatmap illustrating the differentially expressed genes (DEGs; mean expression) based on virushost interactions. Clustering analysis was performed in whole blood samples by age and target using Euclidian distance. B) Venn diagram showing common and exclusive DEGs during aging after matching with SARS-CoV-2-interacting proteins. C) Heatscatter plot presenting 22 DEGs in the whole blood during aging. The color of the circles in the plot reveals the Log_2_FC, while the size reflects the −Log_10_ transformed FDR adjusted p-value. Fold-change was used to represent gene expression in blood samples in the three most advanced ages (50-59, 60-69, and 70-79 years old). D) Tissue-specific gene network of blood proteins predicted to interact with SARS-CoV-2. The network was generated using the HumanBase online tool (https://hb.flatironinstitute.org/). E) PPI interaction of proteins identified as differentially regulated illustrating the top three enriched terms and pathways. Functional interaction analysis was performed with STRING (PPI enrichment p-value < 1.0e-16; minimum confidence score: 0.9). F) Functional enrichment analysis (GO terms) using all DEGs of the blood during the aging process was generated in the Metascape tool (https://metascape.org).

We generated a PPI network for all over-represented targets using the STRING database (enrichment p-value < 1.0e-16, highest confidence of 0.09; Figure 1 E). This analysis highlighted *FASLG, SRC, VCAM1, BAG3*, and *HSPA5* as the main interactions involved in immune response, inflammation, and platelet activation and aggregation; these proteins are directly or indirectly associated with the plasma membrane. To identify top enriched terms, we performed gene ontology (GO) analysis using these targets to unravel aged blood’s functional significance with virus-host PPI. The most relevant processes and molecular function included the immune system process, cellular component organization, biological adhesion, signaling and response to a stimulus, regulation of the biological process, and developmental process (Figure 1 F). Although CTSW and CTSE proteins are related to viral endosomal escape, FASLG, VCAM1, and BAG3 interact with viral Orf8 and protein sars7a (Tables S10 and S11). Also, VCAM1 and BAG3 showed potential interaction with spike glycoprotein, E2 glycoprotein precursors, Excised polyprotein 1..4369 (gene: orf1ab), and Full_polyprotein 1..4382.

### Gender-dependent transcriptional responses in whole blood during aging

We further compared mean gene expression in a cohort with female and male samples to identify gender-dependent responses (GTEx, release V8). When we compared male and female blood samples together, we found that expression of *FASLG, CTSW, CTSE*, and *BAG3* genes significantly increased throughout aging (Figure 2). *VCAM1* was the only transcript that increases its levels in individuals of 60-69 years old compared with younger individuals (20-29 years old). The gene expression profile in the whole blood of male and females samples were similar. Notably, *FASLG* showed an increased expression over the age of 50 in the blood of males and females in a similar manner (Figures S2 and S3). Among all DEGs, *CTSW* presented a similar expression profile in males and females. The *CTSE* was higher in males > 50 years old compared to females. Specifically, *VCAM1* and *BAG3* showed age-dependent expressions when men and women were analyzed individually (Figure S2 and S3).

**Figure 2.**
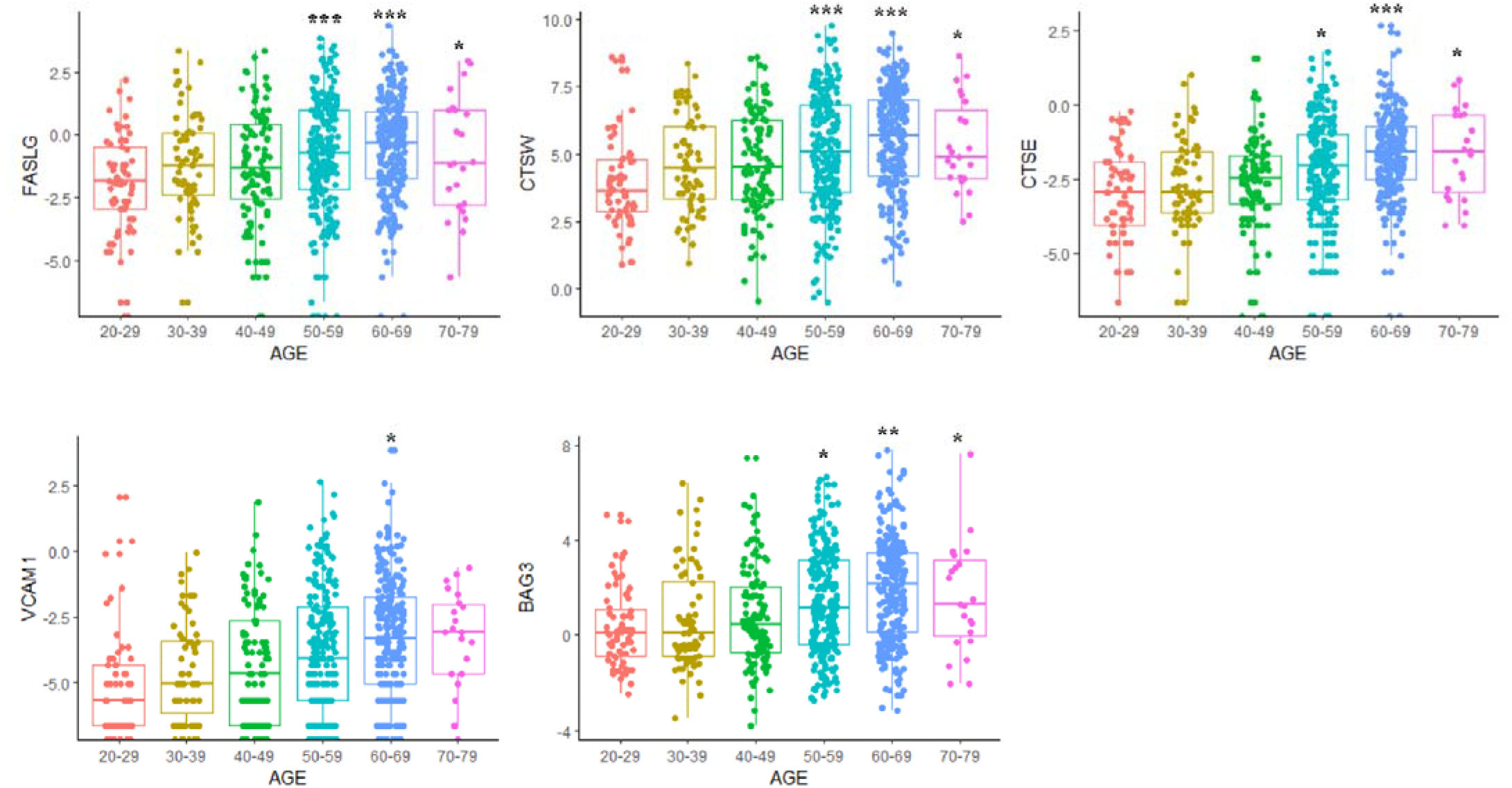
Gene expression levels (TMM) of common targets in whole blood over aging. Data are represented in boxplot by mean ± SD. * P < 0.05, ** P < 0.001, and *** P < 0.0001 vs. young adult individuals (20-29 years old). ANOVA complemented by Tukey’s test. *FASLG*: tumor necrosis factor ligand superfamily member 6; *CTSE*: cathepsin E; *CTSW*: cathepsin W; *VCAM1*: vascular cell adhesion molecule 1; *BAG3*: BAG family molecular chaperone regulator 3.

### FASL is predominantly expressed by NK cells and CD8+ T cells

We analyzed the expression profile of our candidate genes (*FASLG, CTSW, CTSE, VCAM1*, and *BAG3*) to identify blood cell subpopulations in single-cell RNA sequencing (scRNA-seq) data of peripheral mononuclear blood cells from SARS-CoV-2-infected patients. Of note, natural killer (NK) cells, CD8+ T cells, and alpha-beta memory T cells were associated with the expressions of *FASLG* and *CTSW* genes, the latter being expressed by a higher number of immune cells (Figure 3 A and B). Also, *BAG3* gene was mainly expressed in CD4+ alpha-beta memory T cells, naïve T cells, and in CD14+ monocytes (Figure 3 C). We also noticed that *VCAM1* and *CTSE* genes were minimally expressed by these blood cells (Figure 3 D and E).

**Figure 3.**
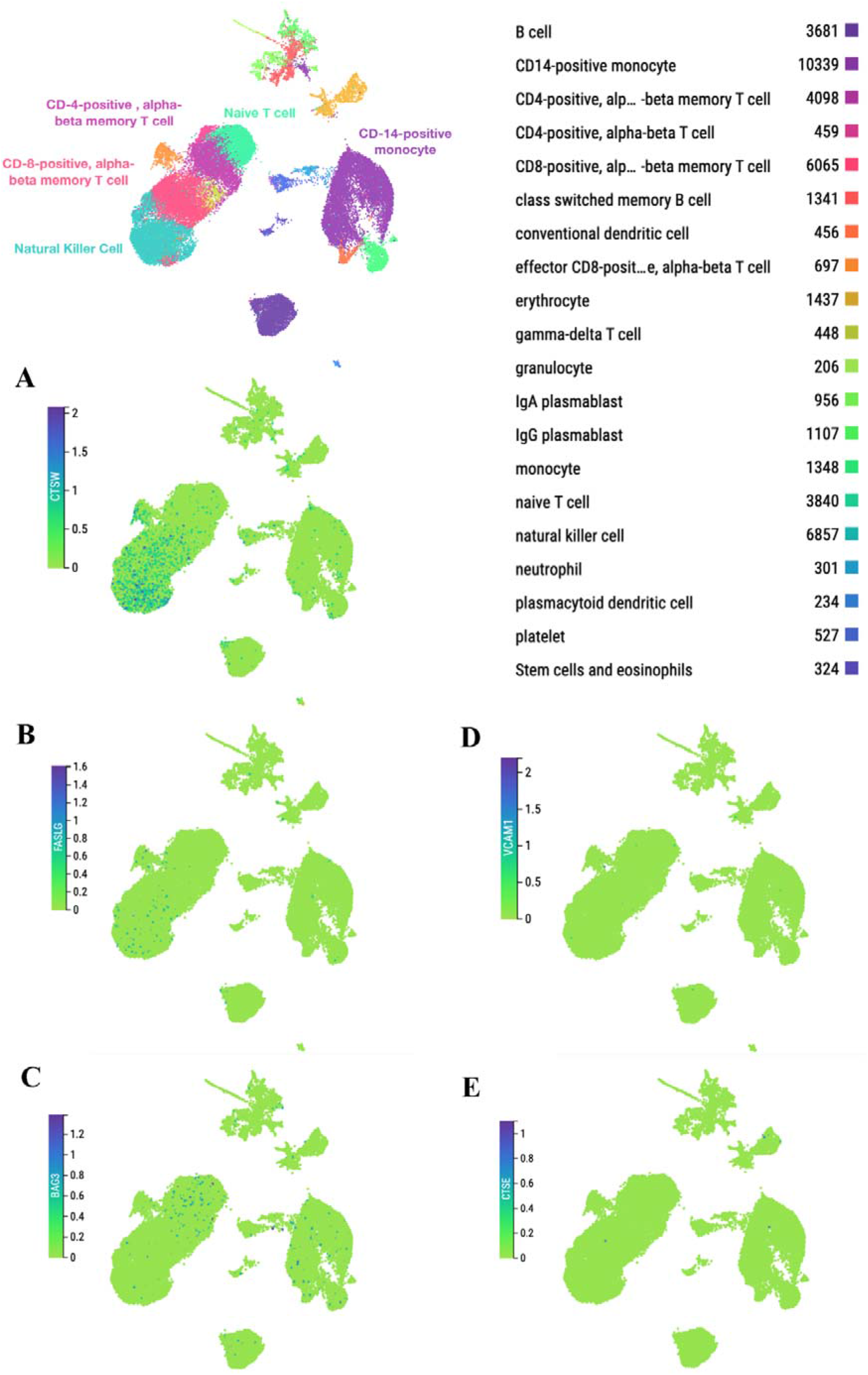
Single-cell gene expression analyses of *FASLG, CTSE, CTSW, VCAM1* and *BAG3* in peripheral mononuclear blood cells (PMBC). Regional clustering depicting distinct cell populations identified in the blood samples of SARS-CoV-2-infected patients in a *t*-distributed Stochastic Neighbor Embedding (tSNE) plot (upper panel). Green dots represent single cells from PMBC samples, and these were not included in the analysis. The range represents the minimum and maximum expression. Colored cell subgroups are shown by the total number in the legend.

Fas ligand (FasL) comprises 281 amino acids (aa) and is divided into a C-terminal intracellular domain (1-80 aa), a transmembrane domain (81-102 aa), a N-terminal extracellular domain (103-281 aa) (Figure 4 A). While FasL N-terminal extracellular domain is responsible for self-association, the FasL C-terminal intracellular domain has a function of receptor binding (PDB ID: 4MSV, chain A; Figure 4 B). Its Fas receptor comprises 335 aa, which are divided into a putative signal sequence of 16 aa (the N-terminal domain with 155 aa, the transmembrane region with 19 aa, and the C-terminal domain with 145 aa (Figure 4 C and D). Although structurally unresolved, the complex Fas/FasL is formed by binding of trimeric FasL to Fas, resulting in signaling-competent trimers that lead to the assembly of death-inducing signaling complex (DISC) with intracellular proteins. Figure 5 summarizes the possible mechanistic role of Fas/FasL signaling activation after SARS-CoV-2 viruses enter the bloodstream and interact with T cells and NK cells. This may result in exacerbated immune response throughout the organism in addition to inducing exhaustion of particular blood cells.

**Figure 4.**
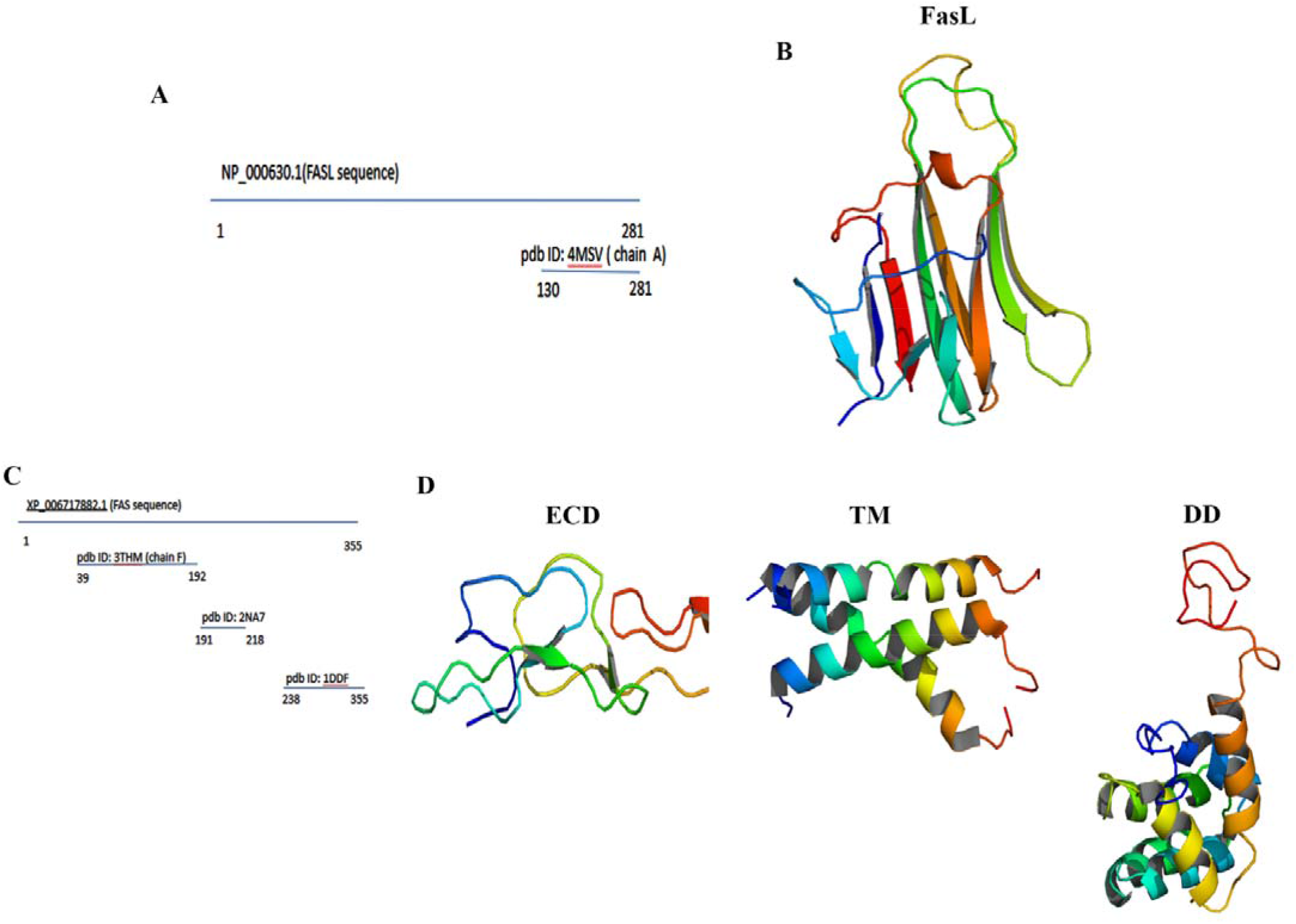
FasL and Fas high resolution structures. A) Alignment of the full FasL sequence with the pdb structures sequences. B) Extracellular domain of the FasL, PDB ID: 4MSV chain A. The structure is shown in cartoon mode from N-terminal (blue color) to C-terminal (red color). C) Alignment of the full Fas sequence with the pdb structures sequences. D) Fas receptor structure displaying the extracellular domain (ECD), PDB ID: 3THM chain F (left panel), the trimer association of the transmembrane domain (TM), PDB ID: 2NA7 (middle panel), and the cytoplasmic death domain (DD), PDB ID: 1DDF (right panel). The structures are shown in cartoon mode from N-terminal (blue color) to C-terminal (red color).

**Figure 5.**
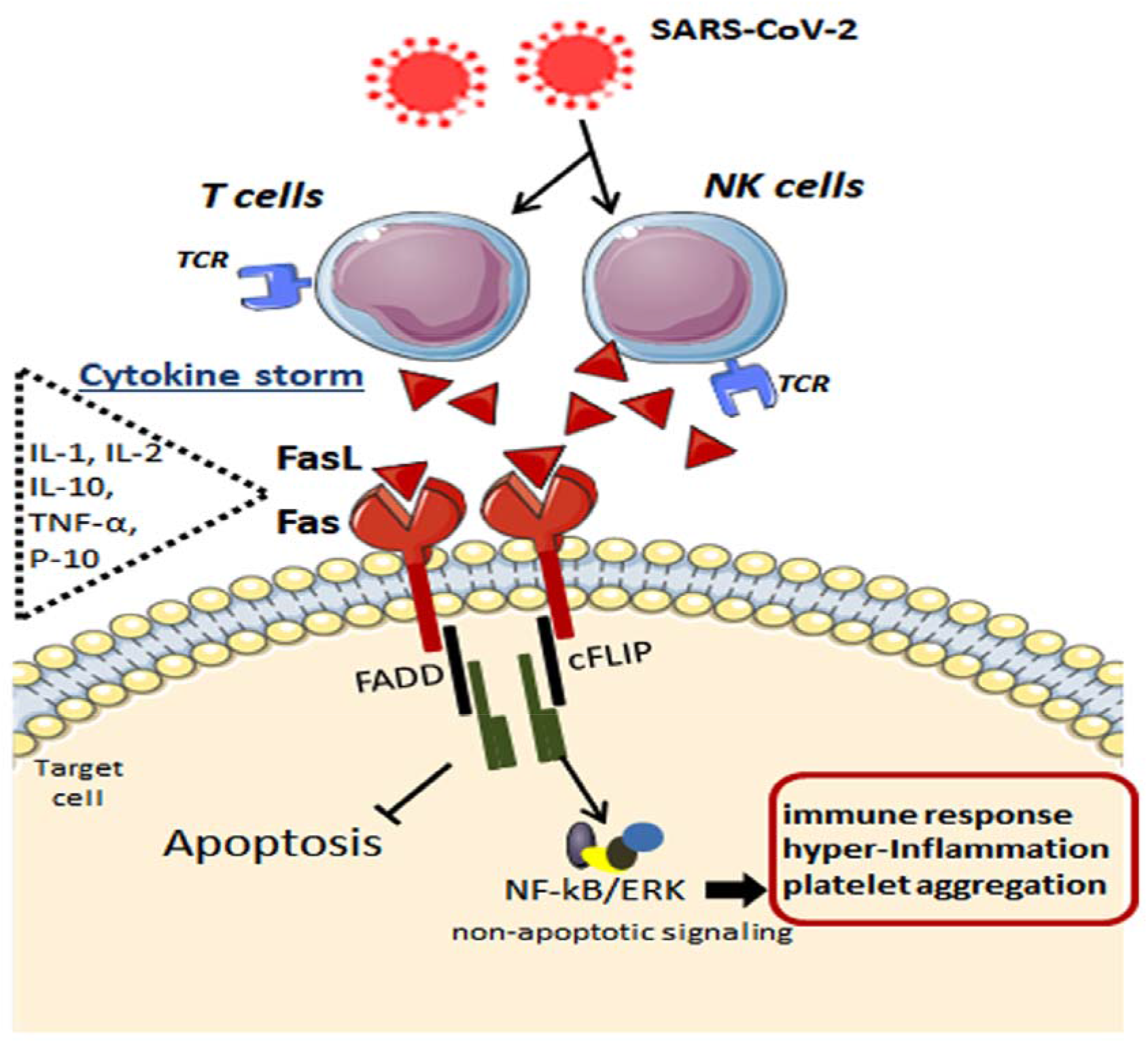
Possible mechanisms underlying the Fas-FasL-mediated signaling in the immune cells after SARS-CoV-2 infection. By infecting specific immune cells, the cytokine storm takes place with elevation in interleukins and chemokines such as IL-1, IL-2, IL-10, IFNs, TNF-α, IP-10, like others. In addition to inflammatory mediators, FasL ligand is highly released by these immune cells, which, in turn, bind to its Fas receptor in the target cell. Upon FasL binding, the apoptosis inhibitor cFLIP is upregulated at the post-translational level and is associated with TRAF1 and TRAF2 and with the kinases RIP and Raf-1, resulting in activation of NF-kB transcription factor and ERK signaling. These anti-apoptotic signals lead to potentiated pro-inflammatory effects via Fas engagement, while FasL increases activated human T cells’ proliferation. Fas, death receptor; FasL, Fas ligand (tumor necrosis factor ligand superfamily member 6); NK, natural killer; T, type of lymphocyte; TCR, T-cell receptor; IL-1, interleukin1; IL-2, interleukin2; IL-10, interleukin 10; TNF-α, tumor necrosis factor alpha; IP-10, interferon gamma-induced protein 10; FADD, FAS-associated protein with death domain; cFLIP, FADD-like IL-1β-converting enzyme-inhibitory protein; NF-kB, nuclear factor kappa B; ERK, extracellular signal-regulated kinase.

## Discussion

The multi-systemic involvement associated with rapid clinical deterioration is among the hallmarks of COVID-19-related mortality. Blood components are deeply involved with widespread virus dissemination and disease aggressiveness; recent approaches uncovered that coronavirus’s immune responses persist beyond six months [33]. To better understand what happens with the blood of patients over aging, we investigated the virus-host interactions using whole blood samples of a young adult compared with older adult individuals. In general, a distinct profile of blood targets was clustered with younger ages (20-49) and older ages (50-79), in which the most distant groups (20-29 years old versus 70-79 years old) displayed an inverse gene expression pattern. The most pronounced effects were observed over the age of 50 and included higher expression of SARS-CoV-2-related genes (e.g., genes involved in immune response, inflammation, cell component and adhesion, biological process, and platelet activation/aggregation). The increase in inflammatory mediators strongly correlates with disease severity within the conception of cytokine storm [34, 35]. Since blood diffuses through the capillaries to guarantee tissue perfusion, the multiple organ dysfunction syndrome may be likely due to essential elements arising from cytokine storm; therefore, monitoring cytokines reduce mortality in other viral diseases such as SARS, MERS, and influenza [36]. Unlike an early infection, the advanced disease is associated with low levels of the antiviral interferons (IFNs) and high levels of interleukin (IL)-1β, IL-6, and tumor necrosis factor (TNF) and chemokines (CCL-2, CCL-3, and CCL-5) secreted by coronavirus-infected immune cells [37, 38].

We provided herein a transcriptomic investigation to identify a possible signature of reliable candidates involved with aged blood. While FASLG was found to be overexpressed in the three highest age ranges of 50-59, 60-69, 70-79 years, some cathepsins (CTSW and CTSE), adhesion-related molecule (VCAM1), and chaperone regulator molecule (BAG3) were commonly increased after the age of 60. FasL is a type-II transmembrane and homotrimeric protein belonging to the tumor necrosis factor (TNF) family After binding to its Fas receptor, a type-I transmembrane TNF receptor, FasL triggers apoptotic and highly inflammatory activities [39]. Although Fas-FasL signaling has shown involvement in apoptosis of immune cells and virus-infected-target cells [40], emerging evidence highlights the apoptosis-independent role of Fas-FasL on the induction of active pro-inflammatory signals in severe pathological conditions (e.g., viral infection) [41, 42]. FasL promotes T cell activation in humans by recruiting cFLIP to the DISC, thereby activating NF-kB and ERK/AP-1 transcription factors [43]. These activations were responsible for the secretion of IL-2 and T cell proliferation; IL-8 was also associated with NF-kB transactivation in bronchiolar epithelial cells, whereas macrophages secreted TNF-α after Fas ligation [44] without triggering apoptotic signaling. Consistently with these findings, we detected by single-cell analysis that FasL is mainly secreted by CD8+ T and NK cells in the peripheral blood of SARS-CoV-2-infected patients. More importantly, a possible interaction of viral Orf8 with a variety of host proteins including FasL may explain the rapid spread of the coronavirus and immune evasion, since Orf8 of SARS-CoV-2 is highly secreted and downregulated MHC-I in cells [45]. A recent study by Sorbera et al. [46] on specific SARS-CoV-2-induced targets reported Fas-FasL signaling involved with endothelial function and neutrophil lifespan and related SARS-CoV-induced apoptosis with potential viral replication. Thus, inhibiting Fas/FasL interaction may be useful in the treatment of COVID-19.

A comprehensive study using whole blood samples from 54 COVID-19 patients documented a dramatic increase in immature neutrophils in parallel with a decrease in CD8+ T and VD2 γδ T-cells count, which is likely due to its differentiation and activation [47]. Based on this fact, we believe that the low count of FasL-associated CD8+ T cells could result from its activation. The immune responses to SARS-CoV-2 infection are often characterized by hyperactivation of CD4+ and CD8+ T cells [48, 49] and macrophages [50], which produce massive levels of pro-inflammatory cytokines. Current evidence reports that critically ill patients have elevation in IL-6 levels compared to moderately ill patients [35]; in addition to the infiltration of inflammatory cells, immune cells have been found in patients’s lung tissues. Notably, disease-associated transcriptional change in aged whole blood had a more pronounced overlap with control blood in comparison to lung tissue transcriptome [51]. By interacting host genes with SARS-CoV-2 and blood transcriptome, Bhattacharyya and Thelma [51] further suggested that viral infection only alters expression profile already dysregulated in the elderly, thereby resulting in poor prognosis; these altered blood genes may reinforce the appearance of severe clinical manifestations including strokes, blood clots, and heart failures.

The expression of CTSW, CTSE, VCAM1, and BAG3 was further shared in the two last age ranges compared to young individuals. Cathepsins SW (CTSW) and SE (CTSE) are papain-like cysteine protease and intracellular aspartic protease, respectively. These molecules were mainly overexpressed in males compared to females. The cathepsins B/L have been described to mediate viral entry into host cells via the endosomal pathway, participating in cell death, protein degradation, autophagy, and immune activities [52–54]. Although CTSB and CTSL are mainly associated with SARS-CoV-2 infection, CTSW is involved in the escape of viral particles from late endosomes during influenza A virus (IAV) replication [55]. Otherwise, CTSE is expressed in immune cells being implicated in antigen processing MHC class II pathway [56]. Targeting CTSW and CTSE may also be a promising alternative to treat COVID-19.

Serum levels of VCAM1 are elevated in mild COVID-19 and highly increased in severe cases [57]. There are several pathological evidence of thromboembolism, diffuse endothelial inflammation, and viral infection of endothelial cells [58, 59] that are strictly related to disease severity and dysfunctional coagulation. Importantly, it is of great value to investigate the expression of endothelial cell adhesion molecules in aged COVID-19 patients.

The BAG3, an anti-apoptotic co-chaperone molecule referred to as BCL2-associated athanogene 3, was upregulated in naive T cells, CD4+ T cells, and CD14+ T cells from aged individuals. BAG3 is involved in a variety of biological processes such as cell survival and apoptosis, cellular stress response, and cell migration, and is suggested to be part of the SARS-CoV machinery for replication [60]. In this context, BAG3 inhibition seems to promote a significant suppresion of viral replication. Like VCAM1, the BAG3 is predicted to interact with spike (S) glycoproteins. These surface molecules favor virus attachment, fusion, and entry into host cells as a direct target involved in immune responses, being lately evaluated for design and development of the S protein-based vaccines [61]. After SARS□CoV□2 enters the bloodstream, a cascade of events occurs resulting in blood clots and strokes [62]. We verified upregulation of proto-oncogene (SRC) and heat-shock protein 70 member 5 (HSPA5) genes in aged individuals (70-79 years old); these are linked to platelet aggregation, activation, and signalings. The other upregulated targets showed involvement in adaptive immunity, immunoglobulin domain, T cell receptor signalings, and cell adhesion. Future approaches are needed to evaluate the role of these blood targets considering COVID-19-related comorbidities and individual physical conditions over aging.

## Conclusion

In summary, we demonstrated that blood gene expression of FASLG, a Fas receptor-ligand triggering non-apoptotic inflammatory activities, is overexpressed in NK cells and CD8+ T cells of males and females after the age of 50. Our in-depth study further evidenced the increase in some cathepsins, cell adhesion, and co-chaperone molecules potentially involved in host cell entry, replication, and vascular dysfunction during aging. Because hypercytokinemia is described as the framework for disease severity and high-risk death, we highlight FASL as a prognostic biomarker and a therapeutic proposal to modulate inflammation in elderly patients with COVID-19. Additional studies are encouraged to test the presence of this biomarker in different disease modalities.

## Supplementary Materials

Supplementary materials can be downloaded directly at the site.

## Conflict of interest

The authors declare no conflicts of interest.

## Funding

The following grant supported the development of this study: São Paulo Research Foundation (FAPESP grant # 2019/00906-6).

## Authorship

LGAC, RFC: conception of the idea, design of the study and drafted the manuscript; JSS: statistical analysis and acquisition of data; MON, MCM: physical and structural analysis and interpretation. All authors significantly contributed with compilation of the literature and approved the final version of the manuscript.

## Supplementary Tables and Supplementary Figure Legends

**Table S1.** Differential gene expression in whole blood transcriptome of individuals aged 20-29 vs 30-39.

**Table S2**. Differential gene expression in whole blood transcriptome of individuals aged 20-29 vs. 40-39.

**Table S3**. Differential gene expression in whole blood transcriptome of individuals aged 20-29 vs. 50-59.

**Table S4.** Differential gene expression in whole blood transcriptome of individuals aged 20-29 vs. 60-69.

**Table S5.** Differential gene expression in whole blood transcriptome of individuals aged 20-29 vs. 70-79.

**Table S6.** Shared upregulated and downregulated DEGs in the whole blood during aging.

**Table S7.** Top 10 enrichment of upregulated shared targets in whole blood during aging.

**Table S8.** Top 10 enrichment of downregulated shared targets in whole blood during aging.

**Table S9**. DEGs in whole blood of aged individuals before and after matching with P-HIPSTer.

**Table S10.** Human-SARS-CoV Interactome based on in silico computational framework P-HIPSTer (http://phipster.org/).

**Table S11.** Interactome of human-SARS-CoV performed in the P-HIPSTer for aged whole blood.

**Figure S1.**
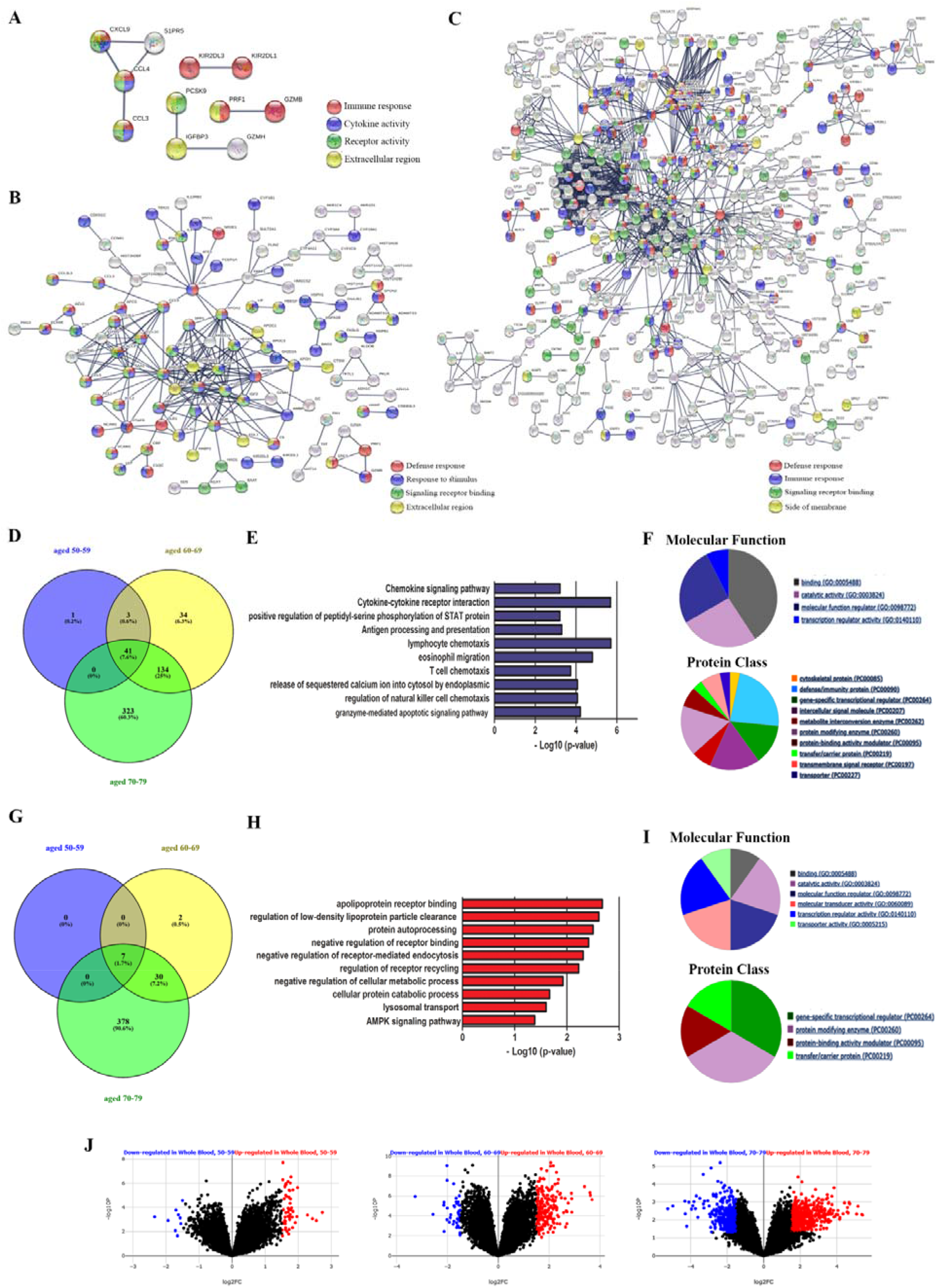
The differentially expressed genes (DEGs) in whole blood were distributed by age to decipher complex networks and molecular activities. A) PPI interaction network (enrichment p-value 0.00441; minimum confidence score: 0.9) of 51 differentially expressed targets in the whole blood of adults aged 50-59. B) PPI interaction network (enrichment p-value value < 1.0e-16; minimum confidence score: 0.9) of 248 differentially expressed targets in the whole blood of adults aged 60-69. C) PPI interaction network (enrichment p-value value < 1.0e-16; minimum confidence score: 0.9) of 908 differentially expressed targets in the whole blood of adults aged 70-79. D) Venn diagram combining blood upregulated transcripts over the age of 50. E) Top functional terms ranking for GO and KEGG pathway categories associated with upregulated shared targets in aged blood identified by gene set enrichment analysis tool EnrichR (p-value ≥ 0.020, combined score ≥ 86.06). F) Pie charts depicting the distribution percentage of molecular functions and protein classes among the blood upregulated molecules (PANTHER classification). G) Venn diagram combining blood downregulated transcripts over the age of 50. H) Top functional terms using GO and KEGG pathway categories associated with downregulated shared targets in aged blood identified by gene set enrichment analysis tool EnrichR (p-value ≥ 0.041, combined score ≥ 75.90). I) Pie charts depicting the distribution percentage of molecular functions and protein classes among the blood downregulated molecules (PANTHER classification). J) Volcano plots showing blood transcriptional gene deregulation in GTEx samples during aging, represented as −log10 (adjusted p-value) and log2 fold change differences. Dashed lines indicate cutoffs (LogFC ≥ |1| ≤ |-1| and FDR<0.05). Blue dots = downregulated targets. Red dots = upregulated targets.

**Figure S2.**
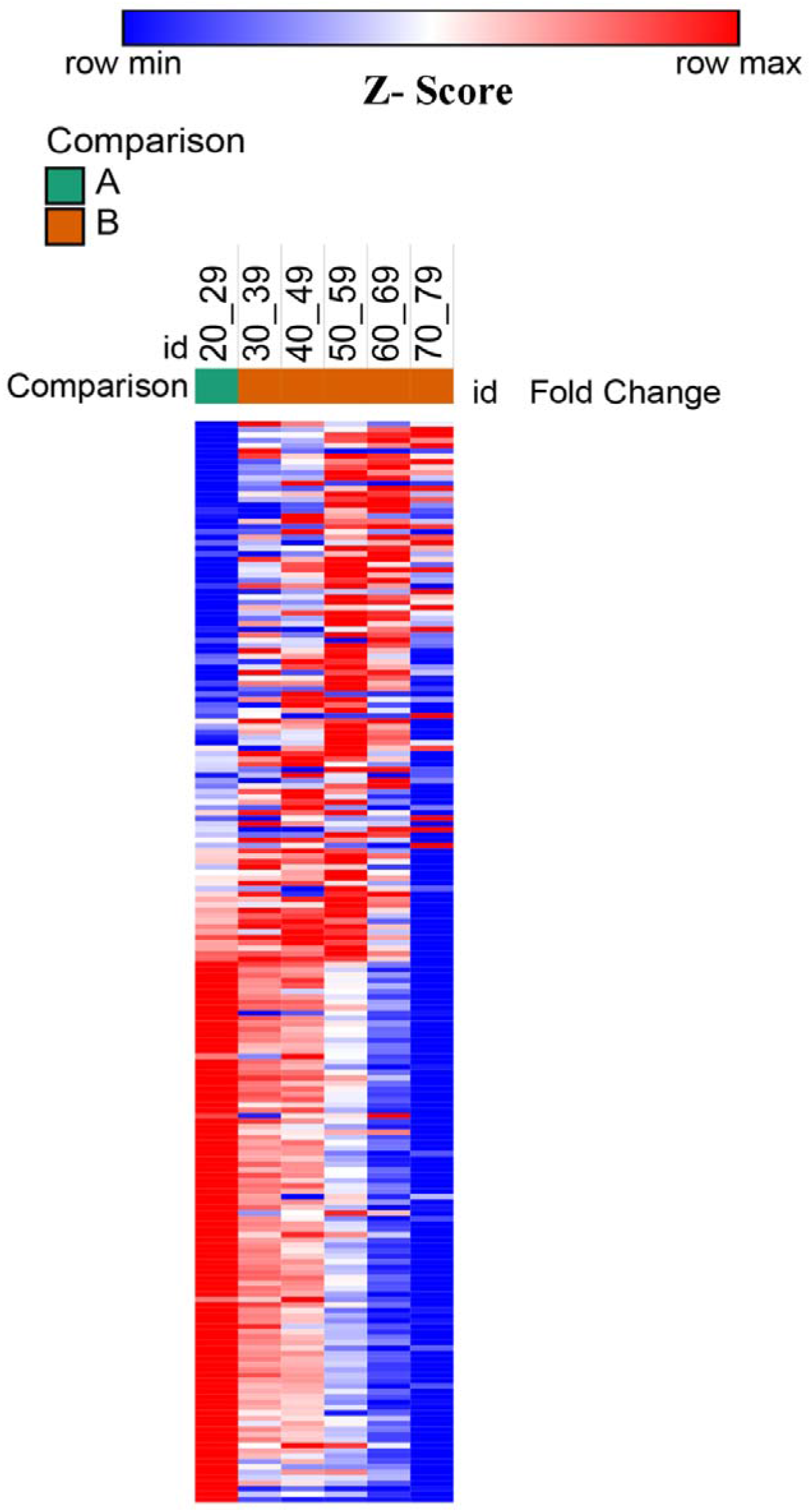
Heatmap of the whole blood samples distributed by age after PPI virushost interaction prediction. Data showing the mean expression of differentially expressed genes, normalized by the trimmed mean of M-values (TMM) and visualized as Z-score. Two factors (A and B) were used for age clustering.

**Figure S3.**
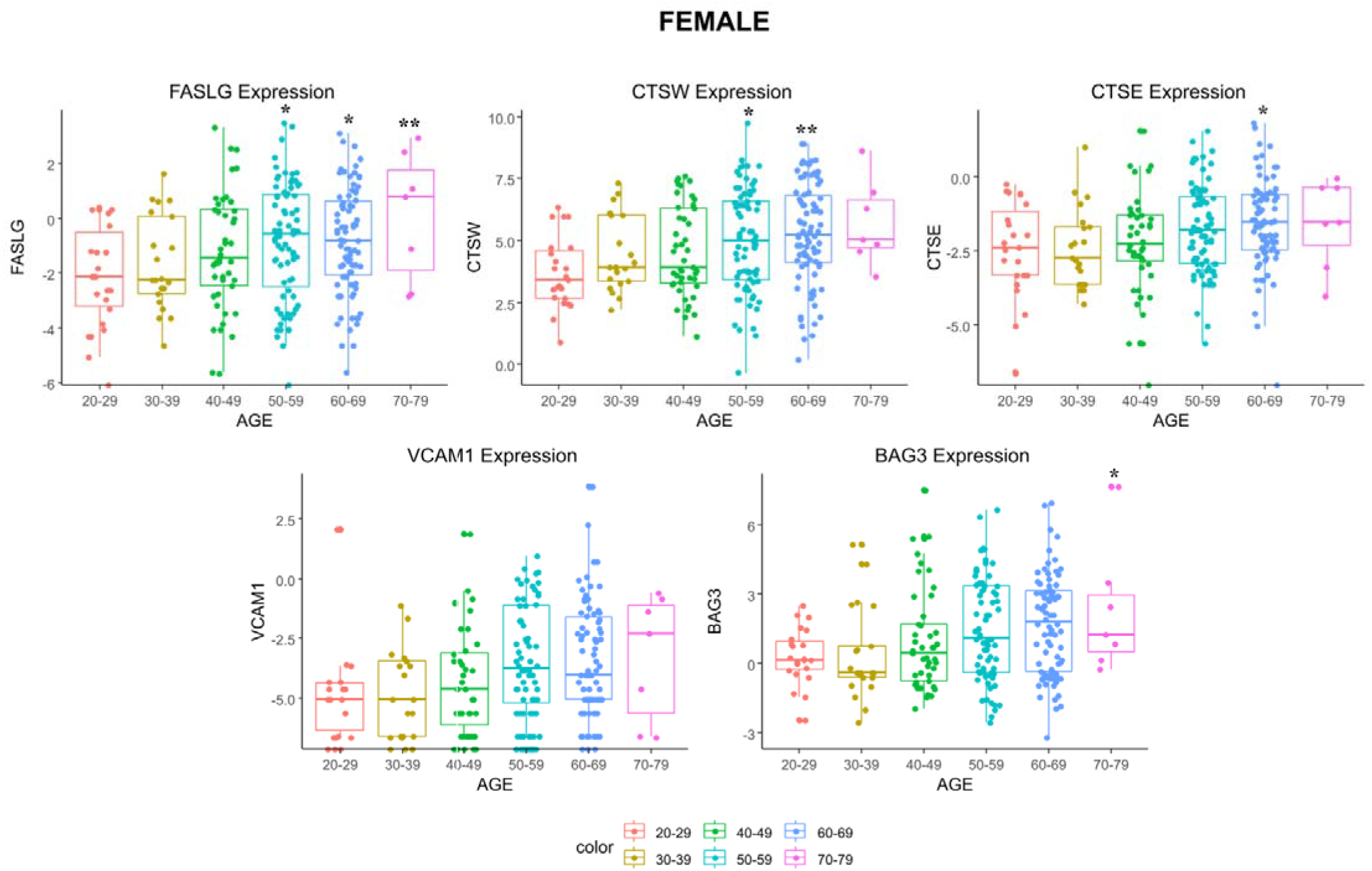
Gene expression levels (TMM) of common targets in female blood samples over aging. Data are represented in boxplot by mean ± SD. * P < 0.05, ** P < 0.001, and *** P < 0.0001 vs. young adult individuals (20-29 years old). ANOVA complemented by Tukey’s test. *FASLG*: tumor necrosis factor ligand superfamily member 6; *CTSE*: cathepsin E; *CTSW*: cathepsin W; *VCAM1*: vascular cell adhesion molecule 1; *BAG3*: BAG family molecular chaperone regulator 3.

**Figure S4.**
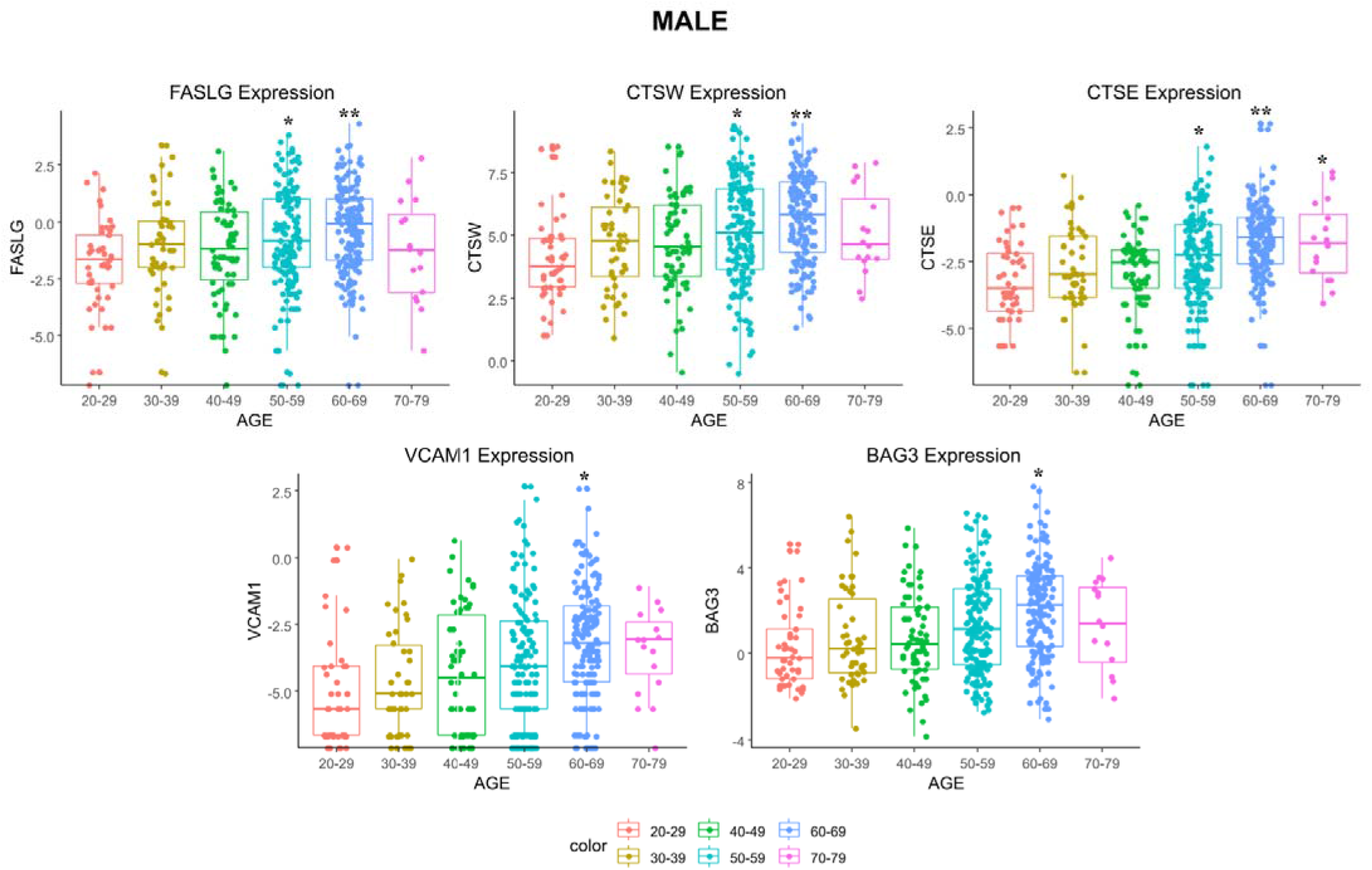
Gene expression levels (TMM) of common targets in male blood samples over aging. Data are represented in boxplot by mean ± SD. * P < 0.05, ** P < 0.001 vs. young adult individuals (20-29 years old). ANOVA complemented by Tukey’s test. *FASLG*: tumor necrosis factor ligand superfamily member 6; *CTSE*: cathepsin E; *CTSW*: cathepsin W; *VCAM1*: vascular cell adhesion molecule 1; *BAG3*: BAG family molecular chaperone regulator 3.

## Notes

### Competing Interest Statement

The authors have declared no competing interest.

